# Reconsidering the border between the visual and posterior parietal cortex of mice

**DOI:** 10.1101/2020.03.24.005462

**Authors:** Sara R. J. Gilissen, Karl Farrow, Vincent Bonin, Lutgarde Arckens

## Abstract

The posterior parietal cortex (PPC) contributes to multisensory and sensory-motor integration, as well as spatial navigation. Based on studies in primates, the PPC is composed of several subdivisions with differing connection patterns, including areas that exhibit retinotopy. In mice the exact anatomical location and composition of the PPC is poorly understood. We present a revised delineation in which we classify the higher-order visual areas RL, AM and MMA as subregions of the mouse PPC. Retrograde and anterograde tracing revealed connectivity, characteristic for primate PPC, with sensory, retrosplenial, orbitofrontal, cingulate and motor cortex, as well as with several thalamic nuclei and the superior colliculus in the mouse. Regarding cortical input, RL receives major input from the somatosensory barrel field, while AM receives more input from the trunk, whereas MMA receives strong inputs from retrosplenial, cingulate and orbitofrontal cortices. These input differences suggest that each new PPC sub-region has a distinct function. Summarized, we put forward a new refined cortical map, including a mouse PPC that contains at least 6 sub-regions, RL, AM, MMA and PtP, MPta, LPta/A. These results will facilitate a more detailed understanding about the role that the PPC and its subdivisions play in multisensory integration-based behavior in mice.

**Highlights:**

- Higher-order visual areas RL, AM and MMA are part of the posterior parietal cortex (PPC) of the mouse based on connectivity.
- The mouse PPC contains at least 6 sub-regions, including RL, AM, MMA, PtP, LPtA/A and MPtA
- Specialized cortical input patterns to the new PPC subdivisions may reflect division of function.
- A new flattened map for mouse cortex represents refined auditory, visual, retrosplenial and PPC areas.

## Introduction

A constant awareness of our posture and position in the world is essential to function adequately in day-to-day life. To enable this, the posterior parietal cortex (PPC) processes information obtained through the different primary senses, and translates it into valid actions and interactions with the environment (Azañón et al., 2010; Bolognini and Maravita, 2007; Mimica et al., 2018; Whitlock, 2017). The PPC is anatomically situated posterior to the primary somatosensory cortex and anterior to the primary visual cortex, integrates different sensory and motor inputs and plays a role in motor, cognitive and spatial navigation-dependent behaviors that make use of egocentric and allocentric reference frames (Chen et al., 2018; Kaas and Stepniewska, 2016; Kastner et al., 2017; Krumin et al., 2017).

Most of the field’s knowledge about the anatomy and function of the PPC comes from research in primates. Anatomically, the primate PPC consists of at least ten cortical regions, including several areas with retinotopic organization. Primate PPC sub-regions display characteristic input and output patterns, especially in their connectivity with other cortical areas, but also with subcortical brain regions. Typical cortico-cortical connectivity involves the sensory and motor areas, such as somatosensory, visual and auditory cortex, but also cingulate, retrosplenial and frontal cortices (Cavada and Goldman-Rakic, 1989a, 1989b; Save and Poucet, 2009; Whitlock, 2017). The higher order lateroposterior (LP) and laterodorsal (LD) nuclei of the thalamus are identified as their subcortical targets (Kamishina et al., 2008; Musil and Olson, 1988; Padberg et al., 2005; Save and Poucet, 2009).

In rats, analogous connectivity has been described for its PPC medial, lateral and posterior sub-regions, with the same cortical areas and the LP, LD and Posterior (Po) nucleus of the thalamus (Chandler et al., 1992; Kolb and Walkey, 1987; McDaniel et al., 1978; Olsen and Witter, 2016; Reep et al., 1994; Torrealba and Valdés, 2008). However, currently there is no evidence that any of the rat PPC sub-regions displays retinotopy.

Based on cytoarchitecture, the mouse PPC is often described as an area consisting of the same three sub-regions as in rat; the medial PPC (mPtA/mPPC), the lateral PPC (LPtA/LPPC) and the posterior PPC (PtP), located between S1 barrel field and secondary visual areas (medial to S1 and anterior to V2 areas) (Hovde et al., 2018; Paxinos and Franklin, 2012). Recently, Hovde et al (2018) has further investigated this cytoarchitectural diversity/organization, adding three higher order visual areas, with PtP partially overlapping with the anterior part of the Rostro-lateral (RL) area and mPtA with the anterior part of the Antero-medial (AM) area, as well as a full incorporation of the Anterior (A) area in LPtA. It remains unconfirmed whether RL and AM are incorporated in the mouse PPC. Both areas, together with A and the Medio-Medial-Anterior (MMA) area, form a string of retinotopically organized visual areas that receive inputs from V1 (Wang and Burkhalter, 2007; Garrett *et al.*, 2014; Zhuang *et al.*, 2017; for review: Glickfeld and Olsen, 2017). Based on the anatomical location of RL, AM and MMA, we hypothesize that they are integral sub-regions of the PPC instead of merely higher order visual areas.

Besides the privileged anatomical location, also functional evidence accumulates and confirms the multisensory nature of the mouse PPC, yet few studies looked at the specific functions of the proposed sub-regions (Guo et al., 2014; Lyamzin and Benucci, 2019; Mohan et al., 2018; Pho et al., 2018). Most of the functional information about RL, AM and MMA comes from studies focused on investigating their visual field coverage or spatial and temporal frequency preferences (Han et al., 2018; Marshel et al., 2011; Zhuang et al., 2017). A couple of recent studies do show that RL gets input from not only visual, but also auditory and somatosensory areas (Gallero-Salas et al., 2020; Olcese et al., 2013).

To expedite research about the computational principles behind multisensory integration and the link with different behaviors in the mouse, knowledge about the precise anatomical location and composition of the PPC is essential.

In this study, we used an anterograde – retrograde tracer approach to characterize the anatomical input and output patterns of RL, AM and MMA. Each area’s connectivity with primary sensory, retrosplenial, frontal and cingulate areas, as well as the thalamus and the superior colliculus, indicates that these areas are integral parts of the mouse PPC. Relative differences in the strength of the cortical inputs to areas RL, AM and MMA seem to predict functional differences for the newly identified sub-regions within the mouse PPC. We integrated all our anatomical data into a new flatmount-type cortical map that describes the novel delineations of the borders between the visual and posterior parietal cortex in the mouse in posterior-anterior and medio-lateral coordinates.

## Material and Methods

### Animals

A total of 23 C57Bl/6J mice were used. All mice originate from Janvier Elevage (Le Genest-St-Isle, France) and were housed in a 10/14-hour dark/light cycle, with access to food and water *ad libitum.* The tracer injections were performed on adult mice (between 120 and 150 days old). All experiments were approved by the Ethical Research Committee of KU Leuven and were in conformity with the European Communities Council Directive of 22 September 2010 (2010/63/EU) and the Belgian legislation (KB of 29 May 2013).

### Viral tracer injections and brain collection

Mice were anesthetized with a mixture of ketamine hydrochloride (i.p., 75 mg/kg, Dechra Veterinary Products, Eurovet) and medetomidine hydrochloride (1 mg/kg Orion Corporation, Janssen Animal Health). Eye ointment (Tobrex, Alcon) was placed on both eyes to prevent ocular dryness. All mice received one cortical injection with either a retrograde modified Herpes Simplex Virus (HSV) tracer hEF1a-TVA950-T2A-rabiesG-IRES-mCherry (Gene Delivery Technology Core, Massachusetts General Hospital) or an anterograde tracer (BDA, 10,000 MW, D-1956, Life Tech, Carlsbad, California). The injection was made in the left hemisphere (300nl, divided in steps of 50nl with 30s in between) at a cortical depth of 450μm. The stereotactic injections were performed in either RL (−2,4; 2,9), AM (−2,4; 2), MMA (−2,4; 1,4) (Fig. 3 and 6) or V1 (−3,2;2,5/-3,5;2.5) (Fig. 1). After the surgery anesthesia was reversed with atipamezol hydrochloride (i.p., 1 mg/kg i.p., Orion Corporation, Elanco Animal Health). Mice were brought back to their home cage, placed on a heating pad, for recovery. One week post-surgery mice were killed with an overdose of natriumpentobarbital (i.p.; 200 mg/ml) in saline (1/3); Vetoquinol N.V./S.A., Aartselaar, Belgium) and post-mortem perfused with 1% paraformaldehyde (PFA) in phosphate buffer (PB, 0.15M, pH 7.42) for 7 minutes. Brains were collected, post fixed in 4%PFA in PB overnight, and stored in phosphate-buffered saline (PBS; 0.1M, pH 7.5) at 4°C until sectioning.

**Figure 1–.**
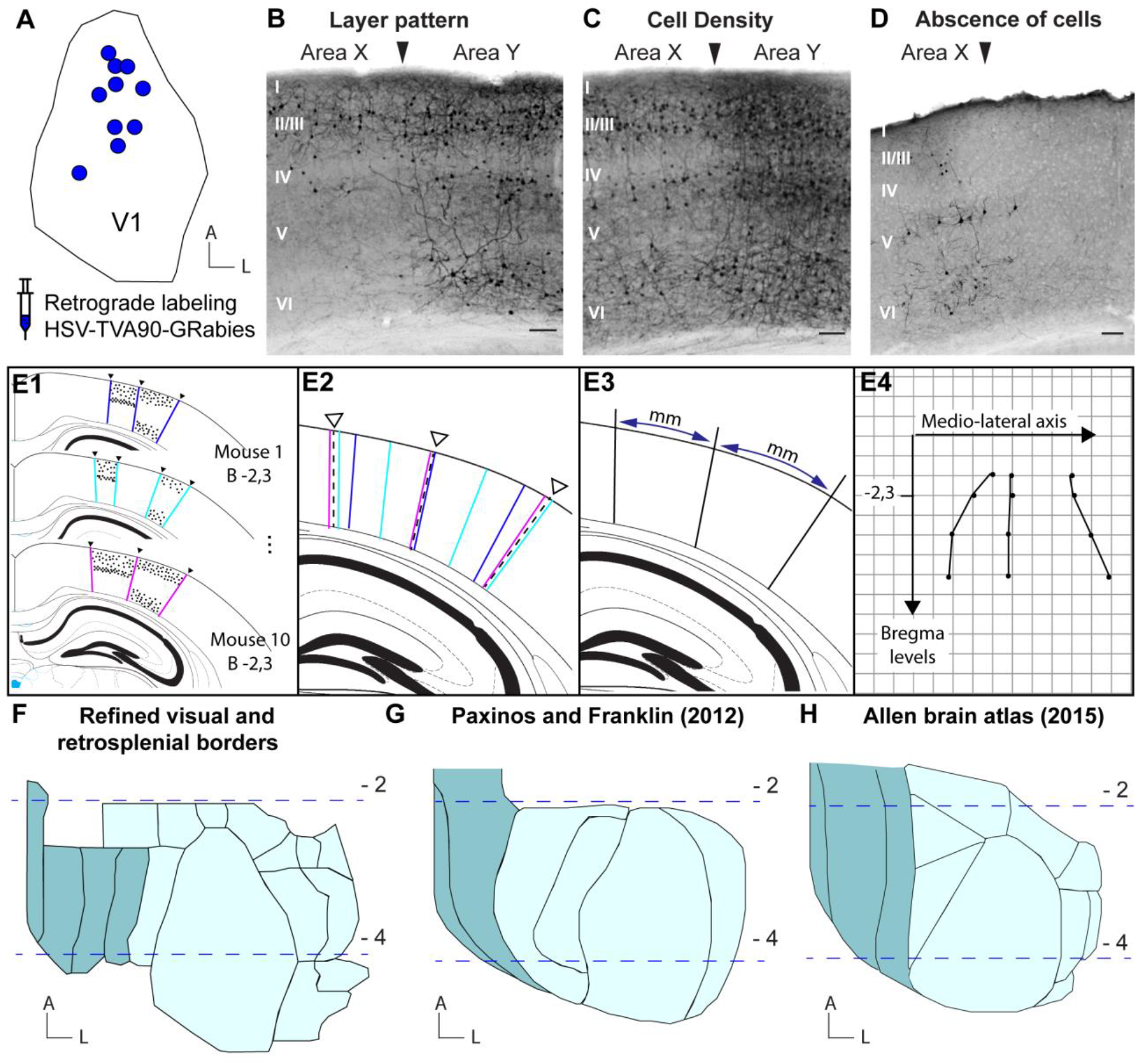
Analyzing and mapping of the borders between visual, posterior parietal and retrosplenial areas in the mouse cortex, based on retrograde tracer injections in V1. **A.** Overview of the 10 injection sites in V1 for the HSV-TVA90-GRabies construct. **B-D.** Illustration of the 3 delineation principles applied: two given areas are labeled as distinct when there is a difference in the distribution of labeled cell bodies across cortical layers **(B)**, a similar labeling across layers but with a different density **(C)** and/or when there is an absence of immunoreactive cells in between two areas with labeling **(D)**. **E1-4.** Schematic representation of the reconstruction of each interareal border, starting from mouse-specific border determination to an averaged flattened map. Borders were delineated on each coronal section from 10 mice **(E1);** and merged for each bregma level on a standard section from the Paxinos and Franklin (2012) atlas **(E2)**. All individual borders were then averaged into one fixed border and the distance between such borders measured while taking the curvature of the cortex into account **(E3).** Every border was then mapped onto a grid with AP and ML coordinates in mm to compile all data, thereby creating the new flattened cortical map **(E4).** **F.** Representation of the final reconstruction of the averaged delineations for retrosplenial (turquoise) and visual cortex (light blue). Dashed blue lines indicate Bregma levels −2 and −4. **G.** Current delineations of retrosplenial (turquoise) and visual cortex (light blue) according to the Paxinos and Franklin (2012) atlas. Dashed blue lines indicate Bregma levels −2 and −4. **H.** Current delineations of retrosplenial (turquoise) and visual cortex (light blue) according to the Allen Brain (2015) atlas. Dashed blue lines indicate Bregma levels −2 and −4.

### Immuno-histology

Coronal sections (50μm-thick) were made on a Vibratome (VT1000S Leica Leitz Instruments, Heidelberg, Germany) and stored in 24 well-plates in PBS. Immunohistochemistry was performed to visualize the injected tracer via light microscopy. Brain slices were incubated for 20min on 37°C and rinsed with Tris-buffered saline (TBS, 0.1M, pH 7.6). After pre-treatment with 0,3% H_2_O_2_ and additional washing steps, they were immersed in 1/5 pre-immune goat serum in Tris-NaCl-blocking buffer (TNB, TBS, 0.1% triton, 0.5% blocking buffer (FP1012, Perkin Elmer, Waltam, Massachusetts, USA) for 45min. Overnight incubation with the primary antibody rabbit-anti-RFP (1:20,000; Rockland 600-401-379, Pennsylvania, USA) was followed by TBS washing steps. Sections were incubated with a secondary antibody Goat-Anti-Rabbit-Biotinylated (1:250 in TBS, 30min) followed by another series of rinses. Streptavidin labeled with Horseradish Peroxidase was applied (1:500 in TBS) for 1 hour. Sections were transferred to 0.6M acetic acid solution (pH 6) after rinsing with TBS and further stained with the glucose oxidase-diaminobenzidine-nickel or SHU method (Shu et al., 1988, Van Der Gucht et al, 2001). This method results in a permanent black staining. Sections were mounted on gelatinized glass slides and dried overnight on a heating plate (37°C). The glass slides were gradually dehydrated via an alcohol series (50% – 70% – 98% – 100% – 100%, 5min each), submerged in Xylol and coverslipped with DePeX. Every section (50μm) was stained and at least every 300μm there was a double stained section for RFP and VGlut2 (1:20,000, Abcam ab79157, Cambridge, UK). Also these sections were stained in series. After performing the anti-RFP SHU staining, the sections were thoroughly washed with PBS (3x 10min) and a second immuno-staining was performed against VGlut2 prior to mounting. Before the DAB staining they were incubated in Tris-stock solution.

### Anatomical mapping of cortical areas

The anatomical mapping of cortical boundaries is based on data from 10 mouse brains and a total of 728 sections (Fig.1, Suppl. Fig.1). Series of sections were either stained for RFP or a combination of RFP and VGlut2. All sections, ranging from bregma −1,9 until bregma −5, were analyzed and photographed with a brightfield microscope (Zeiss Axio, Zeiss, Oberkochen, Germany) at 5x magnification. They were arranged from anterior to posterior and bregma levels were determined by overlaying the sections with those from the Paxinos and Franklin brain atlas (2012). Inter-areal borders were delineated on each section in relation to the immunoreactive cells, by distinguishing groups of cells and their layering pattern (Fig. 1B). For each bregma level (every 100μm) the borders were merged on a standard section from the Paxinos and Franklin, (2012) atlas and averaged into one fixed border. The distance from the midline to a given border was measured, taking the curvature of the brain into account, up until the rhinal fissure. Every border was mapped as a distance in mm onto a grid with AP and ML coordinates, leading to the creation of the flattened cortical map described in the results section (Figure 1). This was done by using Image J and Adobe Illustrator.

### Analysis of the input patterns

Groups of immunoreactive neurons were assigned to specific cortical areas. Defining the cortical areas on coronal sections was based on available atlases (Allen Institute for Brain Science, 2015; Paxinos and Franklin, 2012) and on recognizable landmarks, like the barrels, as visualized via the VGlut2 marker, the presence and appearance of hippocampus and subcortical structures like the thalamic nuclei. On top of that, our own research data and delineation of cortical areas, which lead to the creation of a new cortical flattened map for mouse, helped in the identification of the labeled areas. Every cortical area with immunoreactive neurons was studied over its full anterior to posterior axis. The coronal section with the most immunoreactive neurons for a given area was chosen for analysis. Comparing this type of counting versus counting all the immunoreactive cells within the complete area along the anterior-posterior axis has shown that there is a dismissive difference between the two. For this reason we have chosen to analyze the input patterns to RL, AM and MMA by counting cell bodies on the most labeled section for each cortical region that gives input. The number of cell bodies counted (Fiji, Image J, Cell counter, (Schindelin et al., 2012)) per area was then divided by the total number of counted cell bodies throughout the brain, to calculate the fraction of input from a given cortical area to the injection site.

## Results

To confirm or refute the prediction about the anatomical parcellation of the mouse PPC, afferent and efferent connectivity to a set of higher-visual areas, RL, AM and MMA, was characterized using antero- and retrograde tracing. To facilitate the necessary tracer injections the relative anatomical location of these areas was first determined in brain coordinates in relation to surrounding visual, somatosensory, auditory and retrosplenial cortices and the previously described anterior subregions of the PPC, LPtA, mPtA and PtP.

### Refinement of inter-areal borders for visual and retrosplenial cortex

We determined the exact position and inter-areal borders of RL, AM and MMA in relation to bregma and the midline by visualizing the different areas that give input to V1 via injection of a retrograde tracer in ten mice and by targeting two retinotopic positions within V1 (Fig. 1A). The retrograde-labeled neurons projecting to V1 were detected in series of coronal brain sections by immunostaining against the RFP reporter protein from the HSV tracer. Every group of immune-reactive cells was allocated to a cortical region based on all possible combinations of three principles, layer pattern, cell density and absence of immunoreactive cells (Fig. 1B-D). Figure 1B illustrates a border between two areas based on a layer pattern difference. Area X has immunoreactive cells in the supra- and granular layers, but not in the infragranular layers, while area Y has labeled cells in all cortical layers. The second, cell density principle is illustrated in panel C of figure 1. Area X and Y have immunoreactive cells throughout all layers, but the number of immunoreactive cells in area X is lower than in area Y. The last indicator for an areal border is the absence of cells (Fig. 1D), when immunoreactive cells are only found in area X, and not in the adjacent cortical space. Whereas the occurrence of one of these three principles, layer pattern (Fig 1B), cell density (Fig. 1C) and absence of cells (Fig. 1D), can be sufficient to label two regions as different areas, a combination of multiple principles appeared necessary for other areas. Individual borders were traced on each coronal section, with an interval of 50 μm between Bregma −1.9 and −5, from the midline to the edge of the auditory areas (approximately 0 to −4 ML) (Fig. 1E1). By merging the information of all brain sections, from 10 brains injected in different retinotopic fields (Fig. 1A, Suppl Fig. 1A), we were able to create a complete cortical map, with the help of the above-mentioned principles. Sections of the same Bregma level of each injected mouse were merged on top of the representative standardized section from the Paxinos and Franklin atlas (2012) (Fig. 1E2, Supplementary Fig. 1B-K). The average borders were determined and their distance from the midline was measured, following the curvature of the brain (Fig. 1E3, Suppl. Fig. 1L) to eventually be mapped onto a grid with coordinates (Fig. 1E4). The flattened cortical map in panel 1F illustrates the refined representation of the visual (light blue) and retrosplenial (turquoise) cortex, compared to the Paxinos and Franklin delineations (Fig. 1G) and the Allen brain atlas (2015) (Fig. 1H).

Figure 2 illustrates the delineated areas with labels (Fig. 2A) and different examples of raw data from several mice (Fig. 2B-I). The exact same principles also allowed interpretation of the complete arealization along the anterior-posterior axis of each of the mouse brains (Supplementary figure 2). We successfully delineated areas PM, V1, LM and LI (Fig. 2B). The border between PM and V1 can be identified based on layer pattern. PM has immunoreactive cells in the supragranular layers, but not in the infragranular layers, while V1 has immunoreactive cells throughout all layers. Between V1 and LM there is an absence of cells, with LM and LI also having immunoreactive cells throughout all layers, but with a different cell density. RL, A, AM and MMA can be delineated based on differences in layer pattern and absence of cells (Fig. 2C,D). MMA has a sparser labeling than AM as well as an absence of cells in layers 2/3 and layer 5/6 (Fig. 2C). The absence of cells is more explicit between AM and RL. Moreover, the immunoreactive cells within RL are more densely grouped than those in AM, especially in layer 2/3. Figure 2D shows three areas where A, the middle one, has a very different layer pattern than AM or RL. The number of labeled cells is considerably lower than in AM or RL, across all layers. Delineating LM from LI is based on a different layer pattern and density (Fig. 2E). There is an absence of immunoreactive cells for LI in layer 6, compared to LM, and layer 2/3 of LM shows a much denser labeling than in LI.

**Figure 2–.**
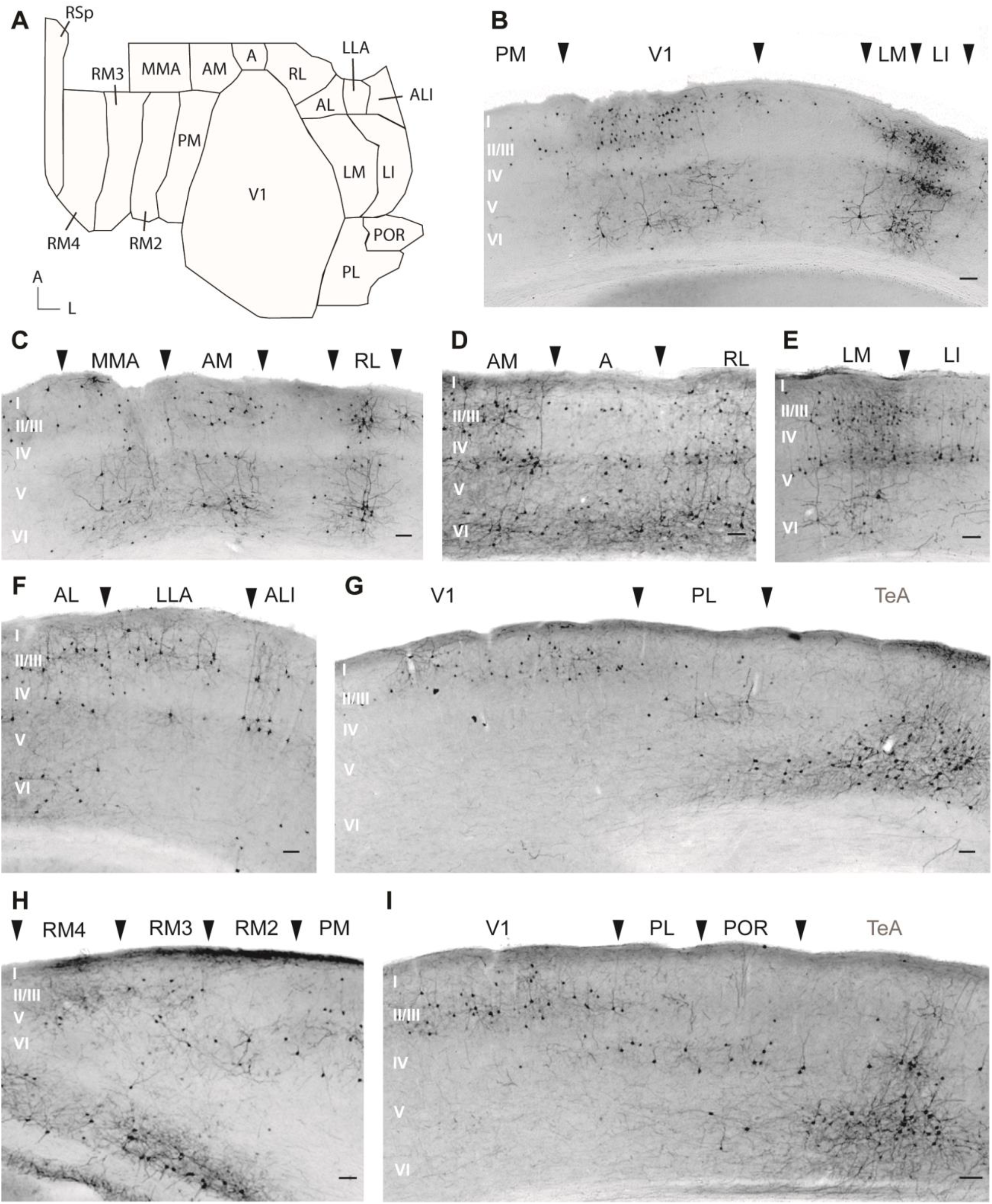
Examples of raw data of mouse specific border determinations. **A.** The new flattened cortical map with the delineated areas denominated. **B-I:** Example coronal sections are shown with black arrows indicating individual areal borders, and related areas denominated. Scale bar = 100μm. See list of abbreviations for clarification of area denominations; Area TeA was not analyzed in detail and is therefore depicted in grey.

Border delineations for AL, LLA and ALI are illustrated in figure 2F. AL has immunoreactive cells throughout all layers, while LLA has mainly immunoreactive cells in the supragranular layers and ALI more from layer 5a. A refined delineation of retrosplenial cortex (Fig. 2H) is based on a distinct layer pattern for PM, RM2, RM3 and RM4. RM4 shows no immunoreactive cells in layer 6, while RM3 has a spread of cells over all layers. This is in contrast with RM2, which shows no labeling in layer 2/3 and is situated in between RM3 and PM, two regions showing labeling throughout all layers. Finally, panels 2G and 2I show a delineation of V1, PL, POR and TeA based on different layer patterns. V1 shows a strong labeling in the supragranular layers, while PL and POR show a reduced or even absence of cells in layer 2/3, but stronger labeling in layer 5a. Also, POR shows labeling in layer 6, and PL does not. A clear border is detected between visual cortex and TeA based on the dense labeling in layers 5 and 6.

This way we were able to create the refined representation of the visual and retrosplenial cortex in a flatmount-type cortical map with anterior-posterior and medio-lateral coordinates to facilitate successful injection of tracers in small cortical areas such as RL, AM and MMA.

### Identification of RL, AM and MMA as PPC subregions

In order to be considered a PPC sub-region a specific set of cortico-cortical and subcortical connections with four groups of predefined anatomical brain regions need to be identified (see Supplementary Figure 3). We screened the mouse brain for exactly these output and input patterns to and from RL, AM and MMA, to reliably designate these areas as PPC sub-regions.

#### Cortical input to RL, AM and MMA

The cortico-cortical connectivity patterns revealed for RL, AM and MMA appeared characteristic for PPC subareas. A retrograde tracer was injected in RL, AM and MMA (Fig. 3A). Afferent inputs from several areas to RL are shown in figure 3. There is clearly the typical input coming from the OFC (Fig. 3B), M2 (Fig. 3C), RSA (Fig. 3D), Auditory (Fig. 3E), Somatosensory (Fig. 3F) and Visual cortex (Figure 3G). The same cortical regions also project to AM and MMA (Suppl. Fig. 4,5).

**Figure 3–.**
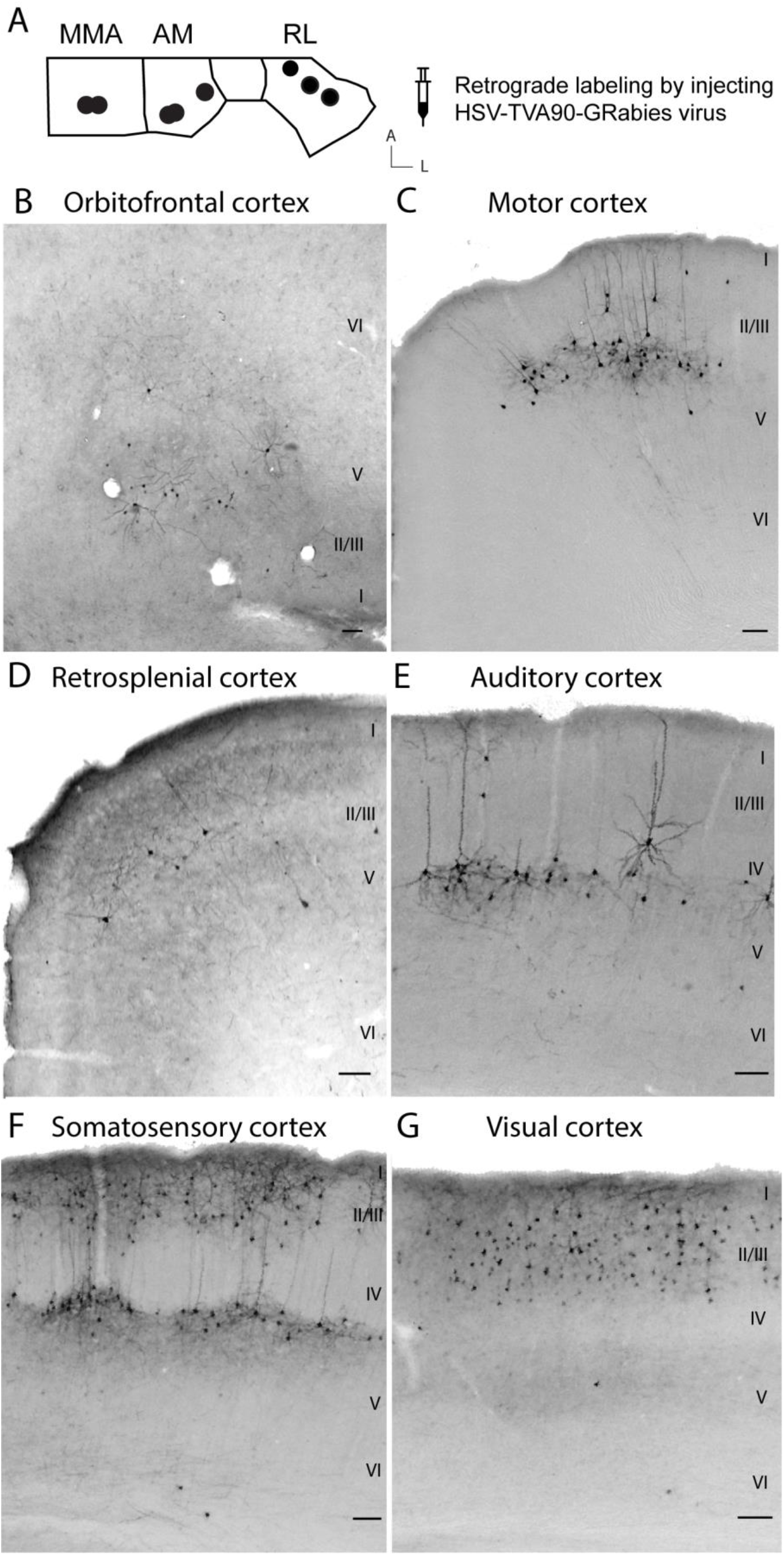
Retrograde labeling of RL shows connectivity coming from all principle cortical areas that qualify RL as a PPC subarea. **A.** Overview of the injection sites for retrograde labeling in RL, AM and MMA. **B-G.** Example neurons projecting to RL were immunolabeled in: **B.** Orbitofrontal cortex; **C.** Secondary motor cortex; **D.** Anterior retrosplenial cortex; **E.** Auditory cortex; **F.** Somatosensory cortex; **G.** Visual cortex. Similar information for AM and MMA is displayed in supplementary figures 5 and 6. Scale bar = 100μm

When comparing the projections to all three areas, clear differences in the relative weight of the inputs to RL, AM and MMA become apparent. Pronounced differences are observed for the motor, retrosplenial, cingulate, somatosensory and orbitofrontal cortices (Fig. 4). Figure 4A summarizes the fraction of relative input percentage from the different motor cortex subregions (M1, M2 and FEF) to RL, AM and MMA. RL receives the most input from the FEF (5.03%), while MMA receives the least (0.59%). M2 gives substantial input to all three areas (between 4 and 6%). Figure 4A2 illustrates the input from the FEF to MMA and Figure 4A3 to RL. We observe a clear difference, with a higher abundance of cells projecting to RL. The same holds for RSA and Cg. Overall, RL and AM receive a very low amount of input from RSA (0.9% for RL and 0.7% for AM) and from Cg (1.22% for RL and 1.72% for AM) (Fig. 4B1). MMA however receives a lot more RSA input (10.4%), specifically from RSAp, as well as Cg input (7.8%; Fig. 4B1). The difference between RSAp input for AM and MMA is illustrated in Figures 4B2 and 4B3. For somatosensory input we found a difference between certain sub-regions and only those have been shown in detail (Fig. 4C1). RL receives the most input from S1BF, more than 20% of its overall input, while MMA barely receives S1BF input (0.1%) (Fig 4C2). AM receives an intermediate amount of S1BF input but does show the most S1T input (4.6%; Fig. 4C1, 4C3). MMA is the only area that receives input from all three OFC sub-regions (Fig. 4D1), mostly from VO (11%), compared to MO (1.7%) and LO (1.06%). AM receives minor input from LO (0.7%) and VO (1.76%), while RL only receives input from LO (1.24%), as illustrated in figures 4D2 and 4D3. Supplementary figure 6 illustrates the extent of variation between brains for input from motor cortex (Suppl. Fig. 6A), retrosplenial and cingulate cortex (Suppl. Fig. 6B), Somatosensory cortex (Suppl. Fig. 6C), and orbitofrontal cortex (Suppl. Fig. 6D). All brains support the same overall conclusion. MMA differs the most from RL and AM, with RL receiving more S1BF input, AM the most S1T, and MMA receiving the most input from RSAp, Cg and OFC.

**Figure 4–.**
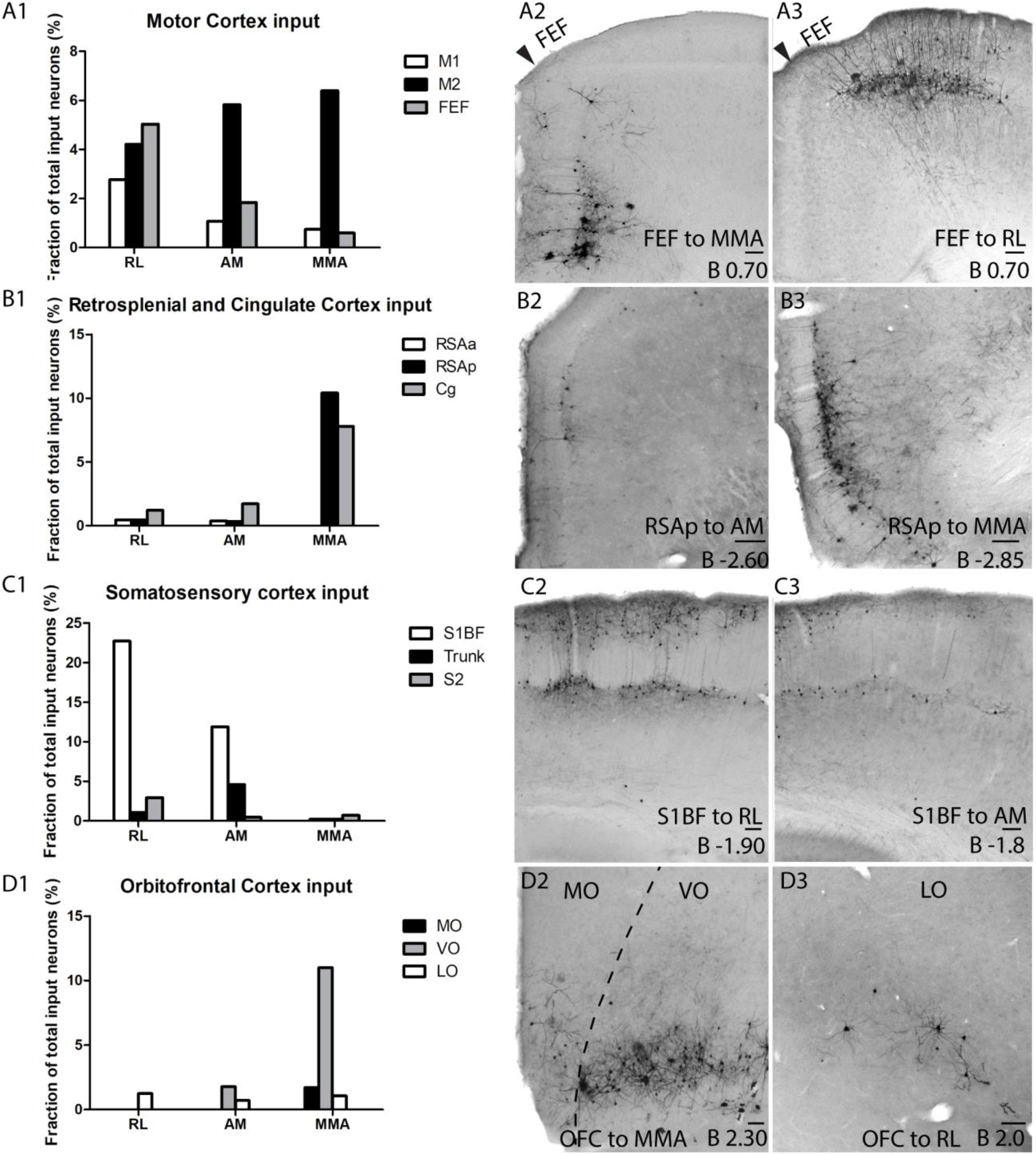
Motor, retrosplenial, cingulate, somatosensory and orbitofrontal inputs to RL, AM and MMA display a different connectional weight. **A1-3.** Input from motor cortex: **A1.** Fraction of total input neurons from M1, M2 and FEF to RL, AM and MMA; illustrated with an example for input from the FEF to MMA **(A2)** and RL **(A3). B1-3.** Input from retrosplenial and cingulate cortex: **B1.** Fraction of total input neurons from RSAa, RSAp and Cg to RL, AM and MMA; illustrated with an example for input from RSAp to AM **(B2)** and MMA **(B3)**. **C1-3.** Input from somatosensory cortex: **C1.** Fraction of total input neurons to RL, AM and MMA; illustrated with an example for input from S1BF to RL **(C2)** and AM **(C3).** **D1-3.** Input from orbitofrontal cortex: **D1.** Fraction of total input neurons from MO, VO and LO to RL, AM and MMA; illustrated with an example for input from MO and VO to MMA **(D2)** and from LO to RL **(D3).**

A schematic representation of the differences in relative input of each cortical area to RL, AM and MMA may facilitate making inferences about functional diversity between the new subdivisions in the mouse PPC. Figure 5 represents the outcome of the quantification of the relative input percentage of the cortical areas to RL, AM and MMA, aligned with a cortical flatmap with similar area-specific color coding. This cortical flatmap merges the existing Paxinos and Franklin atlas (2012), with our refined area delineations (Fig. 1) and info from relevant research papers for the auditory and somatosensory areas (Knutsen et al., 2016; Tsukano et al., 2016). It shows the flattened cortical surface of the left hemisphere of the mouse forebrain, starting from the midline towards the rhinal fissure (X-axis), from bregma 3.2 to −5.2 (Y-axis), with color coding of the different sensory systems (visual (blue), auditory (green) and somatosensory (orange)), motor cortex (red), posterior parietal cortex (purple)) and other cortical areas. By uniting our new and the most recently published delineations of the different cortical areas, this map will facilitate future experimental design, execution and data analysis, by providing clear brain coordinates and revealing details about which areas deserve attention in future research. An enlarged version of the flattened map with brain coordinates can be downloaded from the Supplement (Suppl. Fig. 7).

**Figure 5–.**
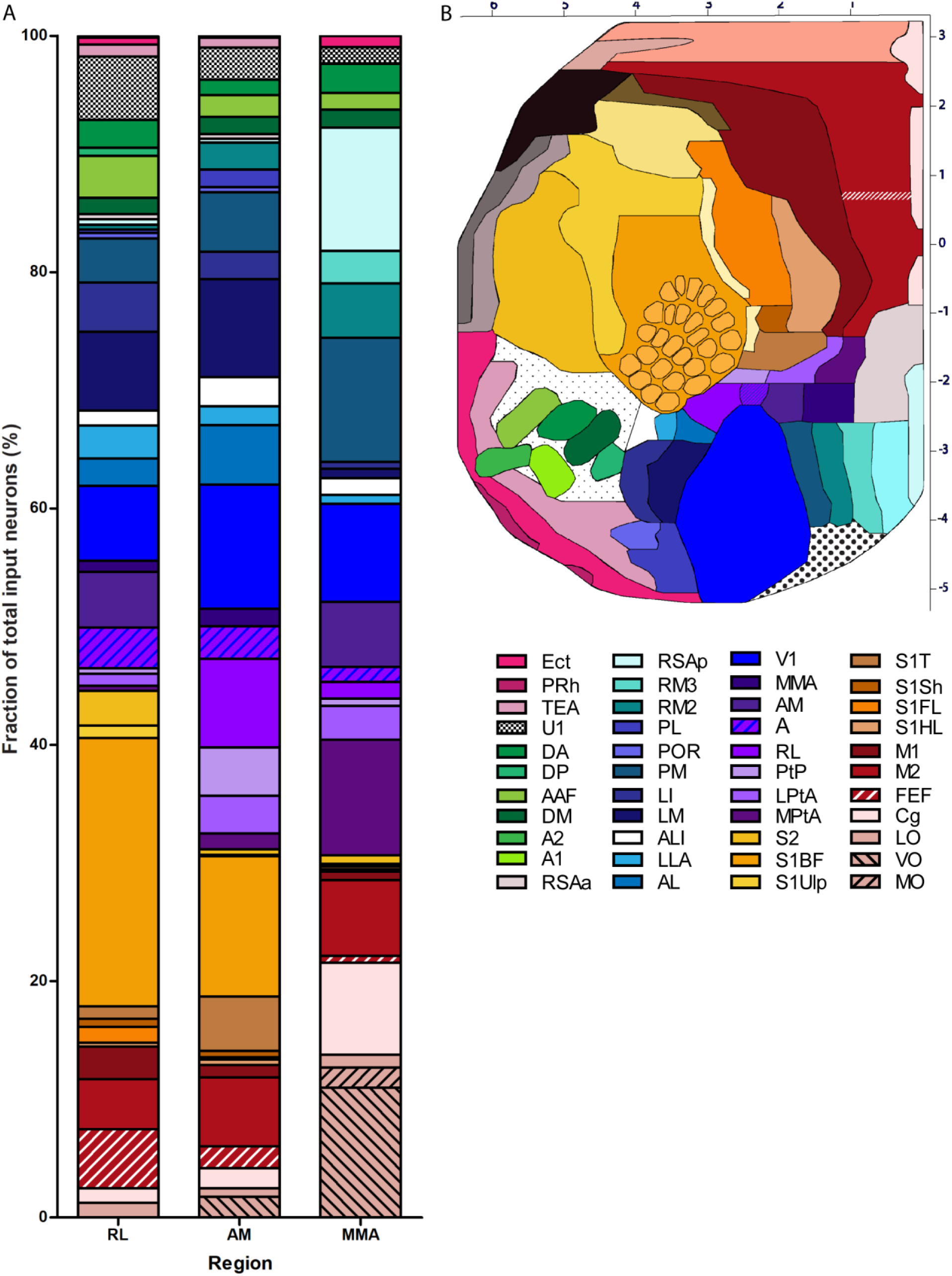
Fraction of cortical input neurons to RL, AM and MMA, from all cortical areas that showed retrograde labeling. **A.** X-axis shows the different areas for which input was analyzed: RL, AM and MMA. The Y-axis shows the fraction of input neurons for every cortical area in which immunoreactive cells were detected. Every area that provides input is color coded, similar to the flattened cortical map **(B)**. **B.** Flattened cortical map with the newly defined areal delineations for visual cortex and PPC (as in Fig. 1F), complemented with the barrel field delineation from (Knutsen et al., 2016) and auditory delineation from (Tsukano et al., 2016), and integrated in the Franklin and Paxinos 2012 atlas. X-axis represents the medio-lateral coordinates in mm, Y-axis represents the anterio-posterior levels in mm. See list of area abbreviations in relation to area denominations.

#### Subcortical output from RL, AM and MMA

Overall, the efferent projections of RL, A and AM to subcortical brain structures confirm that they are part of the PPC and that they do not relay any input to the primary thalamic nuclei. An anterograde tracer, BDA, was injected in RL, AM and MMA to specifically visualize their projections to thalamic nuclei and the superior colliculus. RL, AM and MMA were either injected individually (Fig. 6A, top to bottom) or co-injected with two fluorescent tracers, in RL (green) and AM (red) (Fig. 6E1). Subcortical projections from RL are depicted in Fig. 6 panels B1 to B3, from AM in panels C1 to C3 and from MMA in panels D1 to D3. As predicted, RL (Fig. 6, B1,2 and E3) gives input to the LP (more specifically to LPLR, LPMR and LPMC), LD and Po nucleus, but not to any of the primary sensory nuclei like VPm or LGNd. AM gives input to the same nuclei (Fig. 6 C1,2 and E3), while MMA gives input to the LD nucleus (Fig. 6 D1,2), but not to the Po nucleus and only to the LPMR part of the LP nucleus (Fig. 6 D2). All three areas give input to the intermediate and deeper layers of the superior colliculus, but not to any of the layers above (Fig. 6B3,C3,D3). The input to the intermediate layers is most pronounced. There is a clear medio-lateral topographical component between the input coming from AM and RL to the LP (both to LPLR and LPMR divisions), LD and Po nucleus. This also holds for the antero-posterior axis. This is also the case for the superior colliculus where we observe an areal difference in medio-lateral expression within the layers as well as the antero-posterior axis.

**Figure 6–.**
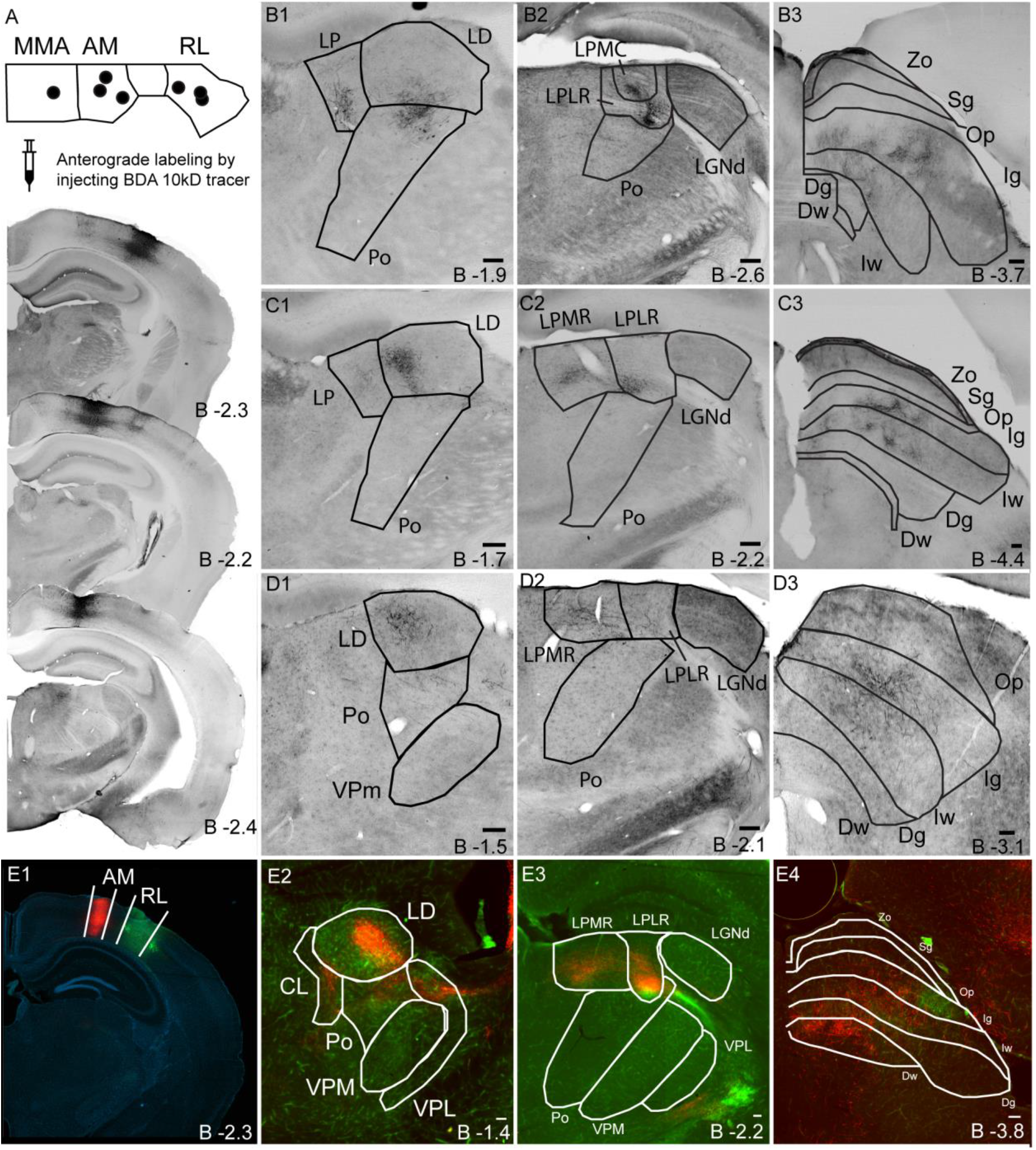
RL, AM and MMA projections to thalamic nuclei and superior colliculus. **A.** Anterograde tracer injections sites for RL, AM and MMA, illustrated by an example for each injected area; from top to bottom: RL, AM and MMA. **B1-3.** Output from RL. **B1 – B2.** Axonal labeling in thalamic nuclei LP (both LPMC as LPLR subdivisions), LD and Po, but not the LGNd. **B3.** Axonal labeling in the intermediate layers (Ig and Iw) of the superior colliculus. **C1-3.** Output from AM. **C1-C2.** Axonal labeling in thalamic nuclei LP (both LPMR as LPLR subdivisions), LD and Po, but not the LGNd. **C3.** Axonal labeling in intermediate (Ig and Iw) as well as deep layers (Dg and Dw) of the superior colliculus. **D1-3.** Output from MMA. **D1-D2**. Axonal labeling in thalamic nuclei LP (Only to LPMR subdivision) and LD, but not the LGNd. **D3.** Axonal labeling in intermediate layers (Ig and Iw) of the superior colliculus. **E1-4.** Co-injections of RL (green) and AM (red) reveal different projection zones to similar target nuclei. **E1.** Injection sites. **E2-E3.** Axonal labeling in thalamic nuclei LD, Po, VPl, CL and LP, but not VPm or LGNd. Both areas project to LPMR and LPLR, but to different (partially overlapping) positions. **E4.** Axonal labeling in the intermediate (Ig and Iw) as well as the deep layers (Dg and Dw) of the superior colliculus. See list of area abbreviations to clarify the area denominations. Scale bar = 100μm.

#### The mouse posterior parietal cortex and its interconnectivity

Our data not only illustrates that the mouse PPC is composed of at least 6 areas, the 3 established areas MPtA, LPtA and PtP and the 3 new additions, RL, AM and MMA, based on cortical input and subcortical output, but also reveals differences in connectivity between these areas. Figure 7 summarizes these findings for the new mouse PPC sub-regions. RL receives the least input from MPtA, LPtA and PtP, compared to AM and MMA, with MMA receiving the most input from MPtA and AM from PtP (Fig. 7A, C). Few cells from RL project to MMA, while its input to AM is much more pronounced. AM on the other hand relays about an equal amount of input to RL and MMA (Fig. 7B, C).

**Figure 7–.**
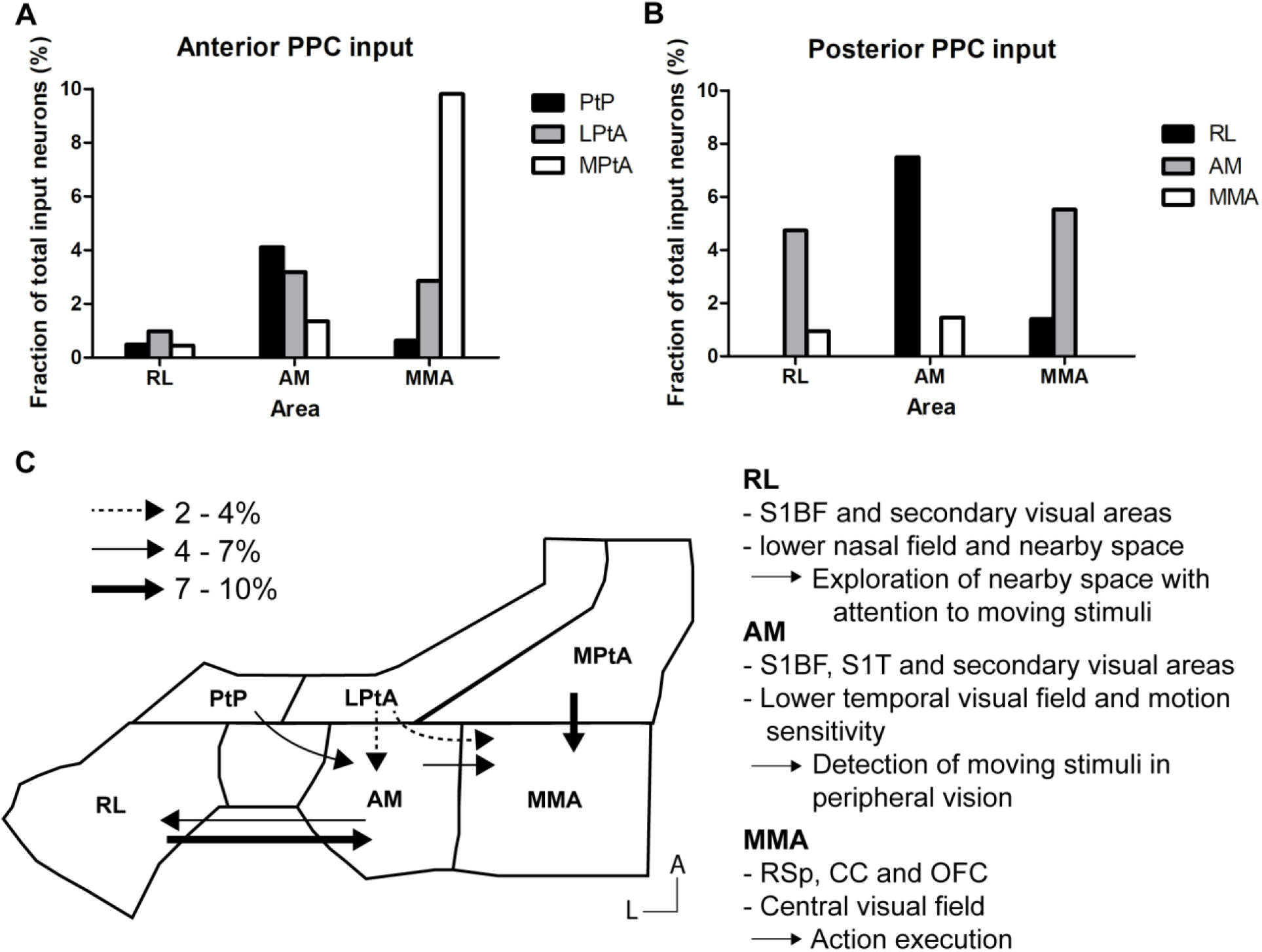
Summary of the findings for mouse PPC subregions and their interconnectivity. **A.** Fraction of total input neurons coming from anterior PPC (PtP, LPtA, MPtA) to subareas RL, AM and MMA. **B.** Fraction of total input neurons between the three new subareas of the posterior PPC (RL, AM and MMA). **C.** Schematic representation of our main findings for the mouse PPC. Different types of arrows represent the fraction of neuronal input towards RL, AM and MMA; Dashed arrows: 2-4%, Regular arrows: 4-7%, Bold arrows: 7-10%. RL receives input from AM, in turn AM receives more input from RL, less from PtP, followed by LPtA; AM gives comparable input to MMA. Besides input from AM, MMA receives more input from MPtA. For each area an additional summary (right) is given of our most pronounced findings related to the cortical connections as established in this paper, and of visual field characteristics from previous research, with a predicted function to instigate new targeted functional studies for mouse PPC subdivisions.

## Discussion

Our data support the presence of least 6 subdivisions in the mouse PPC, including the retinotopically organized areas RL, AM, MMA, and LPtA/A, MPtA and PtP. The three newly added areas, RL, AM and MMA, display the typical connectivity as previously established for primate PPC subdivisions. Areal differences in connectional strength predict differences in multisensory processing. Our anatomical investigations also provide a new flattened cortical map for the mouse, aligned with mediolateral and anteroposterior coordinates (Fig. 5, Suppl. Fig. 7). This knowledge paves the way for new research into a better understanding and a more detailed analysis of current and future mouse PPC data.

### A new flattened cortical map: mouse visual, auditory, somatosensory, posterior parietal and retrosplenial areas in 2D coordinates

We created a new flattened and coordinate-based topview map of the mouse brain. The cortical map merges our new detailed anatomical data for visual, posterior parietal and retrosplenial cortical territory with data from the Paxinos and Franklin (2012) atlas, and from complementary research papers for auditory and somatosensory cortex (Knutsen et al., 2016; Tsukano et al., 2015). The new flat map is in line with the 3D brain from the Allen brain institute, with four significant differences. We revealed a more detailed representation of the mouse PPC, consisting of at least 6 sub-regions. We integrated areas LLA and MMA, regions that had been observed in Imaging based maps of cortical retinotopy (Zhuang et al., 2017), as well as ALI. According to the Paxinos and Franklin atlas, ALI belongs to the visual area of the mouse brain, but so far there were no observations of any retinotopy. Based on its position, it is likely that this area is area 2 (Hishida et al., 2014a; Wekselblatt et al., 2016). If this is the case, ALI/area 2 is more likely driven by all sensory systems instead of being dominated by one. Area 2 receives connections from all three cortical senses, and shows higher activity when there is simultaneous presentation of all three types of sensory stimuli (Hishida et al., 2014b). Since area 2 is often driven directly by the higher order cortical areas, but not exclusively, this area has been linked to identifying the behavioral relevance of a stimulus (Geissler and Ehret, 2004; Wekselblatt et al., 2016), matching different sensory maps (Brett-Green et al., 2003; Yoshitake et al., 2013), as well as translating sensory information to executive areas (Hishida et al., 2014b). The third important addition is the delineation of the retrosplenial cortex into RSAa, RSAp, RM4, RM3 and RM2. This delineation was already described in a previous paper from our research lab based on neurofilament stainings (Van Der Gucht et al., 2007), and is now confirmed based on specific connectional input patterns to V1. Although the neurofilament patterns indicated that RM2 could be a visual area, retinotopy has yet to be found in this area. As a final update, we integrated detailed area information for the auditory system in correlation with the other brain regions. This detailed information (Tsukano et al., 2015) is for now still missing in both the Paxinos and Franklin atlas (2012) as well as the Allen brain atlas (2015).

### RL, AM and MMA are part of the mouse PPC

As an integral part of the PPC, an area needs to connect with a distinctive set of cortical and subcortical brain regions (Whitlock, 2017; Hovde *et al.*, 2018). A PPC sub-region at least connects to the sensory cortices, retrosplenial, orbitofrontal, cingulate and motor cortex, as well as subcortical structures like the thalamic higher order nuclei and the superior colliculus. Based on retrograde and anterograde tracing, we show evidence for such connections for the three higher visual areas, RL, AM and MMA.

All three areas send axons to the superior colliculus, and mainly project to its intermediate layers and in a lesser manner to the deep layers. These projection patterns to the superior colliculus have already been described in detail (Wang and Burkhalter, 2013). The thalamic output connectivity is limited to the LD nucleus, the visual LP nucleus, and the somatosensory PO nucleus. All three areas project to the LD and LP nuclei. The LD nucleus is a higher order nucleus but its precise function remains unclear. There have been connectional studies that link its output to the limbic system, including Cg, and RSP (Spiro et al., 1980), and the LD is possibly involved in spatial orientation and learning, based on somatosensory cues (Bezdudnaya and Keller, 2008; Mizumori and Williams, 1993; Robertson et al., 1983). The LP inputs to visual cortical areas are topographically organized (Juavinett et al., 2019). We observed a similar topography for cortical output to LP, making the topography reciprocal in nature. Both RL and AM project to the somatosensory PO nucleus, supporting a role for these areas in multisensory processing. The PO nucleus can be divided in 4 subareas, POa, POm medial, POm lateral and POT, each receiving projections from the barrels (Sumser et al., 2017). This organization within the PO nucleus is based on projections from S1BF L5b neurons. Projections from AM are limited to the dorsal anterior tip of the PO nucleus (POa), while RL projects to its medial part (POm). The axonal endings are topographically corresponding to the larger straddler whiskers and row 1 and 2. These whiskers are located in the posterior medial S1BF, which corresponds to the observation that most direct S1BF connections to RL and AM come from this S1BF sub-region. The specificity of PPC projections to the POa and POm nuclei could indicate specialized functions within this nucleus corresponding to different PPC sub areas, either related with feedback to S1BF, or to the feed-forward influencing of a subcortical sensory-motor pathway (Sumser et al., 2017). The lack of input to the PO nucleus from area MMA is consistent with the low amount of connections coming from S1 to MMA. This does not however cast doubt on its involvement as a PPC sub-region, since there is no clear evidence that a PPC sub-region should project to both somatosensory and visual higher order thalamic nuclei.

### RL, AM and MMA have distinct connectional patterns

RL, AM and MMA show incoming connections from approximately the same areas, but in different proportions. When looking at the different sensory systems one important noticeable overall difference is that S1 and V1 have direct connections to the PPC sub-regions while only the secondary, and not the primary, auditory cortex makes direct connections. The reason for this remains unclear for now.

One of the most pronounced differences between the PPC sub-regions relates to S1 input. RL receives more than 20% of its connectional input from the barrel field, versus only 12.3% for AM and 2% for MMA. This observation is being backed up by recent imaging studies that observed activity in RL upon tactile engagement (Gallero-Salas et al., 2020; Olcese et al., 2013). Although we did not observe a notable difference in the amount of neurons projecting from V1, it has recently been put forward that RL receives the most projections from V1, compared to AM or MMA (Froudarakis et al., 2019). Retinotopy in RL, as established in previous imaging studies, mainly covers the lower nasal visual field (Garrett et al., 2014; Zhuang et al., 2017). This coincides with current literature pushing RL forward as the hub for connecting spatial and visual input in nearby space (La Chioma et al., 2019; Lippert et al., 2013; Olcese et al., 2013). The preference of RL neurons for high temporal frequency and global motion perception supports an attention to moving stimuli (Han et al., 2018; Juavinett and Callaway, 2015; Marshel et al., 2011). Additionally, we observed that RL receives the highest number of direct projections from M1 and FEF, suggesting a role in visuospatial attention and visual search (Lane et al., 2012).

AM receives less connections from S1BF than RL, yet shows a higher input from the somatosensory trunk area. This might indicate a similar involvement in visual-spatial integration, less dedicated to nearby space but to body-in-space, coinciding with the fact that AM covers the lower temporal visual field (Garrett et al., 2014; Zhuang et al., 2017). Lesions in rat AM have shown that there is performance impairment for visuospatial discrimination (PINTOHAMUY et al., 1987; Sánchez et al., 1997), but not for a visual discrimination task. Inactivating AM in mice during a visual evidence accumulation task, induced an ipsilateral bias, showing a role for AM in making contralateral choices. Inactivating one hemisphere lead to movements in the opposite direction, suggesting a mechanism of spatial hemineglect (Odoemene et al., 2018). AM may thus be responsible for bringing together spatial and visual information from peripheral space. Similar to RL, AM has a high direction selectivity, predicting a function in responding to moving stimuli in the peripheral field (Han et al., 2018; Marshel et al., 2011). Altogether this fits with the extensive input we observed from the trunk and the secondary motor cortex.

MMA covers a much smaller area of the visual field, just below the visual center, covering the same extent of temporal space as AM, but less elaborate in altitude (Zhuang et al., 2017). What role MMA plays remains uncertain due to a sparse amount of research on this area. We observe a significant amount of input from RSAp, as well as from OFC and Cg. This might indicate the involvement of MMA in a sensory-frontal pathway, starting with sensory input activating area 2/ALI and the medial PPC, leading to activity propagation towards frontal areas (Hishida et al., 2014b; Wekselblatt et al., 2016). We found direct connectional input from area 2/ALI to MMA and AM, with AM displaying the strongest connectivity out of all three PPC areas under study as well as a strong connectivity with MMA (Figure 7C). This is suggestive of a circuit where area 2/ALI projects directly to MMA, or indirectly via AM, and subsequently to the Cg and frontal areas. In rat, it was illustrated that the brain areas, M2, Cg, OFC and PPC, show reciprocal connections with each other in a topographic manner (Bedwell et al., 2014; Olsen et al., 2019). This topographic organization has been extensively described for mPPC, lPPC and PtP in the rat brain, showing a medial to lateral gradient in connectivity. In the rat, mPPC is the only PPC subarea that receives connections from all orbitofrontal areas, MO, VO and VLO. The more lateral areas lPPC and PtP only receive projections from VLO (Olsen et al., 2019). We observed a similar topography for the new PPC subdivisions in the mouse, with the most medial area (MMA), being the only one receiving projections from MO. Based on primate research, it is hypothesized that the lateral half of the OFC is involved in identifying the outcome of an individual choice, while the medial OFC compares these different choices to one another, based on a common universal value, leading to influencing what action should be executed (Olsen et al., 2019; Rudebeck and Murray, 2014, 2011; Rushworth et al., 2011). This is further confirmed in rats where MO inactivation showed an increase in choosing the riskier option (Stopper et al., 2014), a shift to reliability on the value of observable outcomes, not predicted ones (Bradfield et al., 2015), and a decrease in probabilistic discrimination (Dalton et al., 2016; Izquierdo, 2017). Another recent observation to support MMA’s function in relation to action execution is that there are multiple cells within the mouse mPPC that respond to different actions (nosepoking, grasping, eating, grooming, turning clockwise, turning counter clockwise and rearing) (Tombaz et al., 2019) and the finding of posture encoding in the mPPC of rats, with higher activity for uncommon postures (Mimica et al., 2018).

## Conclusion

In sum, we demonstrate that RL, AM and MMA are all part of the PPC and propose a mouse PPC that anatomically consists of at least 6 sub-regions. The delineation of all 6 PPC areas resulted in a new flattened cortical map, aligned with mediolateral and anteroposterior coordinates. We not only establish RL, AM and MMA as integral part of the PPC, but also that they have distinct connectional input and output patterns. We propose a role for RL in somatosensory – visual processing of nasal, nearby space and a role for AM in processing somatosensory – visual information in temporal lower space, both with an attentiveness for moving stimuli. On the contrary, we envisage MMA to play a more important role in sensory-frontal/motor processing in guiding the execution of actions (motor) in accordance with the expected outcomes (OFC) and attention to what is behaviorally relevant (area 2/ALI).

## List of abbreviations

A: Anterior area
A1: Primary auditory area
A2: Secondary auditory area
AAF: Anterior auditory field
AL: Anterolateral area
ALI: Anterior Laterointermediate area
AM: Anteromedial area
Cg: Cingulate area
DA: Dorsoanterior field
Dg: Deep gray layer of the Superior Colliculus
DI: Dysgranular insular cortex
DM: Dorsomedial field
DP: Dorsoposterior field
Dw: Deep white layer of the Superior Collliculus
Ect: Ectorhinal cortex
FEF: Frontal Eye Field
FR3: Frontal cortex, area 3
FrA: Frontal Association cortex
GI: Granular insular cortex
Ig: Intermediate gray layer of the Superior Colliculus
Iw: Intermediate white layer of the Superior Colliculus
LD: Laterodorsal thalamic nucleus
LGNd: dorsal Lateral Geniculate Nucleus
LI: Laterointermediate area
LLA: Laterolateral anterior area
LM: Lateromedial area
LO: Lateral orbital cortex
LP: Lateral Posterior thalamic nucleus
LPLR: Lateral Posterior thalamic nucleus, Laterorostral
LPMC: Lateral Posterior thalamic nucleus, Mediocaudal
LPMR: Lateral Posterior thalamic nucleus, Mediorostral
LPtA: Lateral Parietal association cortex
M1: Primary motor cortex
M2: Secundary motor cortex
MMA: Medio medial anterior cortex
MO: Medial orbital cortex
mPtA: Medial posterior associtation cortex
OFC: Orbitofrontal cortex
Op: Optic nerve layer of the Superior Colliculus
PL: Posterolateral area
PM: Posteromedial area
Po: Posterior thalamic nuclear group
POR: Postrhinal area
PPC: Posterior Parietal Cortex
PRh: Perirhinal cortex
PtP: Posterior Parietal association cortex
RL: Rostrolateral area
RM2: Rostromedial area 2
RM3: Rostromedial area 3
RM4: Rostromedial area 4
RSA: Retrosplenial agranular cortex
RSAa: Anterior Retrosplenial agranular cortex
RSAp: Posterior Retrosplenial agranular cortex
S1BF: Primary Somatosensory cortex Barrel Field
S1FL: Primary Somatosensory cortex Front limb
S1HL: Primary Somatosensory cortex Hind Limb
S1Sh: Primary Somatosensory cortex Shoulder
S1Ulp: Primary Somatosensory cortex Upper Lip
S2: Secundary Somatosensory cortex
Sg: Superficial gray layer of the supperior colliculus
TEA: Temporal Association cortex
U1: Unknown area 1
U2: Unknown area 2
V1: Primary Visual cortex
VO: Ventral Orbital cortex
VPL: Ventral Posterolateral thalamic nucleus
VPM: Ventral Posteromedial thalamic nucleus
Zo: Zonal layer of superior colliculus
PtA: Parietal association region

## Acknowledgements

This work was supported by the KU Leuven Research Council (C14/16/048) and the Research Foundation Flanders (FWO), Belgium via research project funding (G061216N) and a doctoral fellowship to S. Gilissen (SB/151597). The funding sources had no involvement in study design, collection, analyses and interpretation of the data or in writing this paper. We would like to thank dr. Nathalie Lombaert, dr. Joao Couto, drs. Maroussia Hennes and drs. Jolien Van Houcke for their helpful comments and suggestions.

## Authors contributions

Study concept and design: SG, VB, KF and LA*. Study supervision: LA*. Data gathering: SG. Analysis and interpretation of data: SG and LA*. Drafting of the manuscript: SG and LA*. Critical revision of the manuscript: SG, VB, KF and LA*. All authors read and approved the final manuscript.

## Data Statements

All data generated in this study can be found in the paper, supplemental materials or can be provided upon request.

