## Supplementary figures for "Reconsidering the border between the visual and posterior parietal cortex of mice"

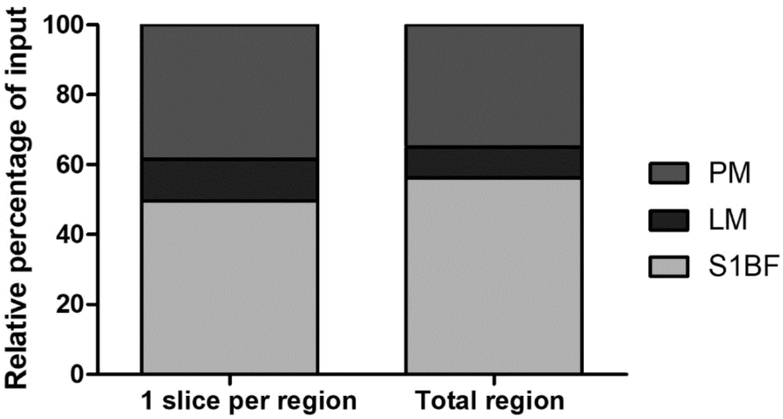


**Supplementary Figure 1 – Counting one coronal section vs counting an entire immunopositive region.** Selection of the coronal section with the most labeling throughout the A-P axis for counting the number of neurons that project to a certain area versus counting all labeled neurons for each projecting area, results in similar findings for the relative numbers of projecting cells, and thus projection weight. There is a dismissible difference between both counting methods. This was tested for two secondary visual areas, areas LM and PM, as well as for the primary somatosensory barrel field (S1BF).

21

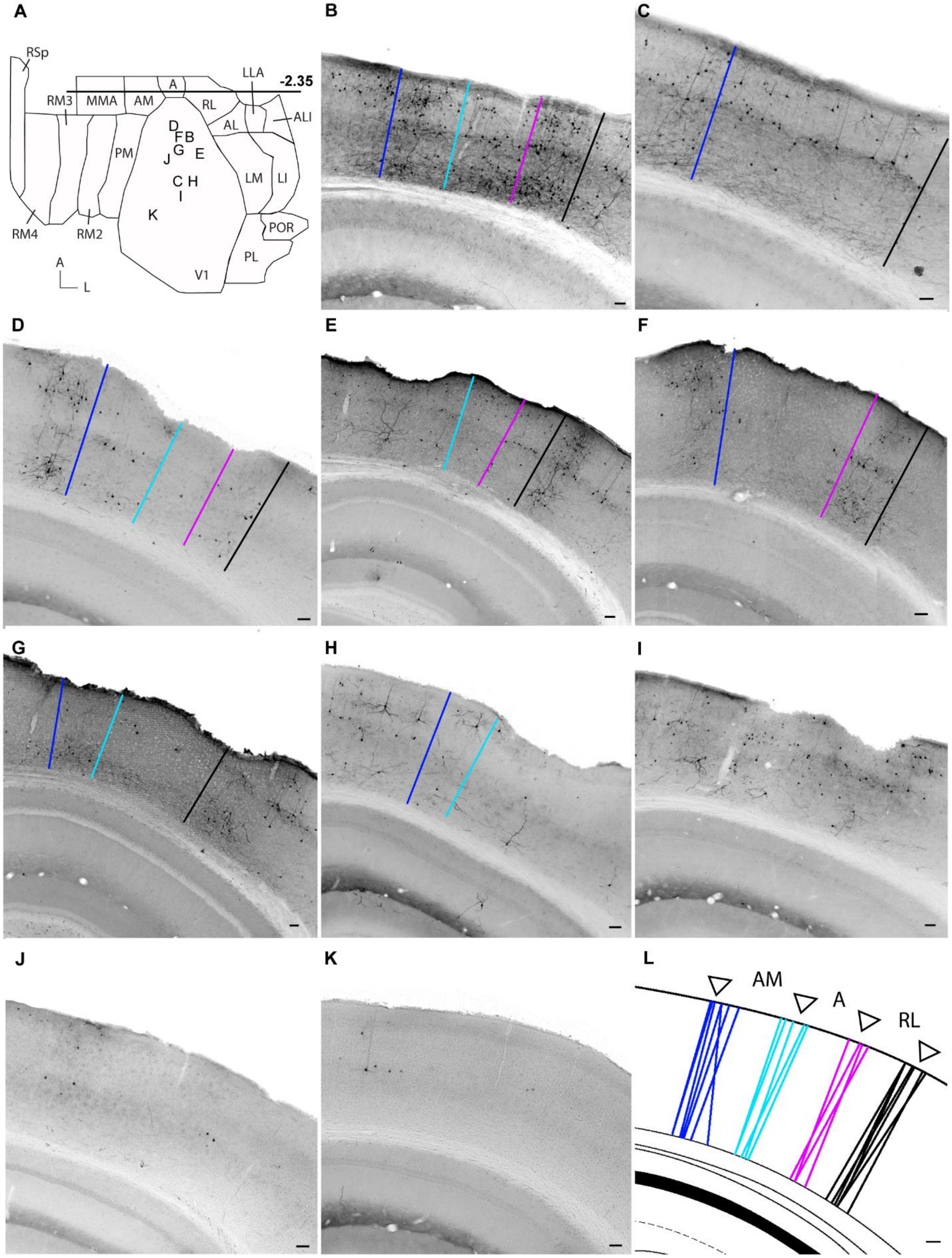

A
B
C
RSp
LLA
-2.35
A
RM3
MMA
AM
RL
ALI
AL
D
F B
G
E
J
PM
C H
I
LM
LI
K
RM4
RM2
POR
PL
V1
A
L
D
E
F
G
H
I
J
K
L
AM
A
RL

**Supplementary Figure 2 – Example of inter-mouse variability in position of similar areal borders when determining the average borders for RL, AM and MMA at bregma -2.35.**

**A.** The new flattened cortical map with the delineated areas denominated. In V1 each letter corresponds to a panel (B-K) and depicts the position of the injection site of the individual examples.

**B-K.** Pictures of raw data at bregma -2.35 for the 10 individual brains. Each color illustrates one border with dark blue = border between MMA and AM, light blue = border between AM and A, pink = border between A and RL and black = border between RL and S1BF. **L.** Summary of all the individual borders, indicating the average border with a white arrow. Scale bar = 100µm.

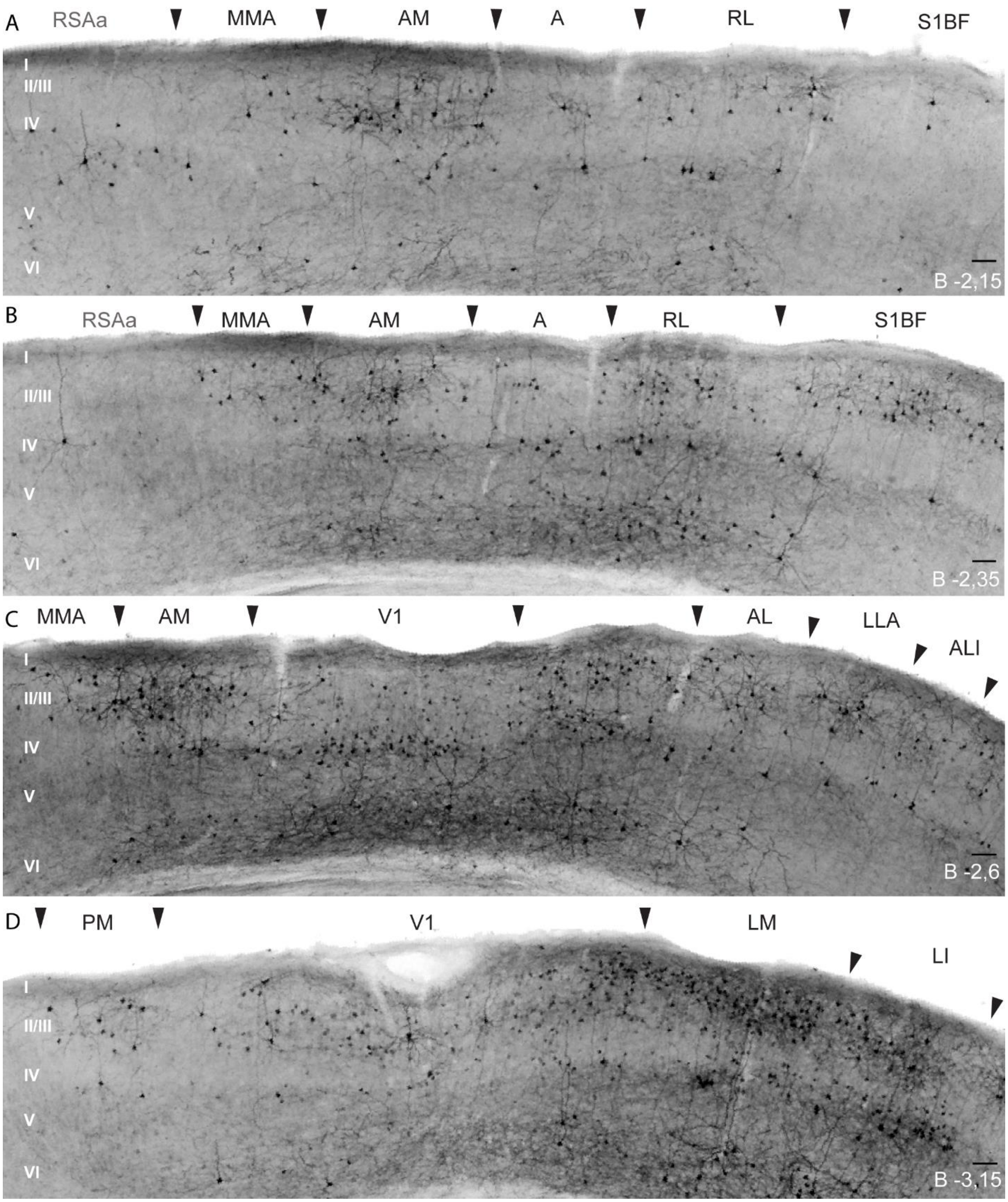


**Supplementary Figure 3 – Illustration of border determination within one brain, on four consecutive sections, from bregma -2.15 to -3.15.** Delineation of RSAa, MMA, AM, A, RL, S1BF, AL, LLA, ALI, V1, PM, LM and LI on coronal brain slices at bregma levels -2,15 **(A)**, -

2,35 **(B)**, -2,6 **(C)** and -3,15 **(D)**. Different layer patterns and density discriminate the different areas, with MMA having almost no IR cells in the lower layers and sparse labeling in layer 2/3, compared to AM which has a much denser labeling in the equivalent layer. AM, A and RL show labeling in all layers, but the amount of cells is less abundant in A than in AM or RL. These area-specific patterns can be observed throughout the brain, as exemplified in supplementary figure 2B, which shows the same areas but 200 µm more posterior in the brain. Overall the density of cells increases but the discriminating factor for A remains the same, with A having a less dense pattern compared to the neighboring areas. When comparing AM and RL, a similar layer pattern is complemented by a more dense label in layer 2/3 compared to 5/6 for AM, while the pattern in RL is more evenly dense throughout the layers. MMA shows the same pattern as in suppl. figure 2A and this analysis can be extended to suppl. figure 2C, which is again 250 µm more posterior, representing the border of these higher order visual regions with V1. A delineation of higher order areas AL, LLA and ALI is shown in suppl. figure 2C. AL has IR cells throughout all layers while LLA and ALI have an absence of cells in layer 5 and 6. They show a similar pattern with a dense clustering of cells in layer 2/3. A delineation of higher order areas PM, LM and LI is shown in suppl. figure 2D. PM shows labeling of cells in the supragranular layers, but not in infragranular layers, while LM and LI mainly show a difference in density and are much stronger labeled compared to PM.

A

**Anatomical groups**

| Subcortical | Sensory | Association | Other |
|---|---|---|---|
| Thalamus | Somatosensory | Orbitofrontal | Motor |
| Striatum | Visual | retrosplenial | Pre-subiculum |
| | Auditory | postrhinal | superior colliculus |
| | | | cerebellum |

B

**Direction of information**

| Give input | Receive input | Reciprocal |
|---|---|---|
| Somatosensory | Superior colliculus | Visual |
| Auditory | Postrhinal | Striatum |
| Cerebellum | Pre-subiculum | Retrosplenial |
| | | Thalamus |
| | | Orbitofrontal |
| | | Motor |

**Supplementary Figure 4 - Connections typifying the mammalian posterior parietal cortex.** **A.** Areas classified based on four anatomical subgroups; Sensory cortex, subcortical, association cortex and other. **B.** Areas classified based on communication direction; Giving input to the PPC (retrograde), receiving input from the PPC (anterograde) or reciprocal. Scheme is based on Whitlock (2017) and Save and Poucet (2009) (Save and Poucet, 2009; Whitlock, 2017).

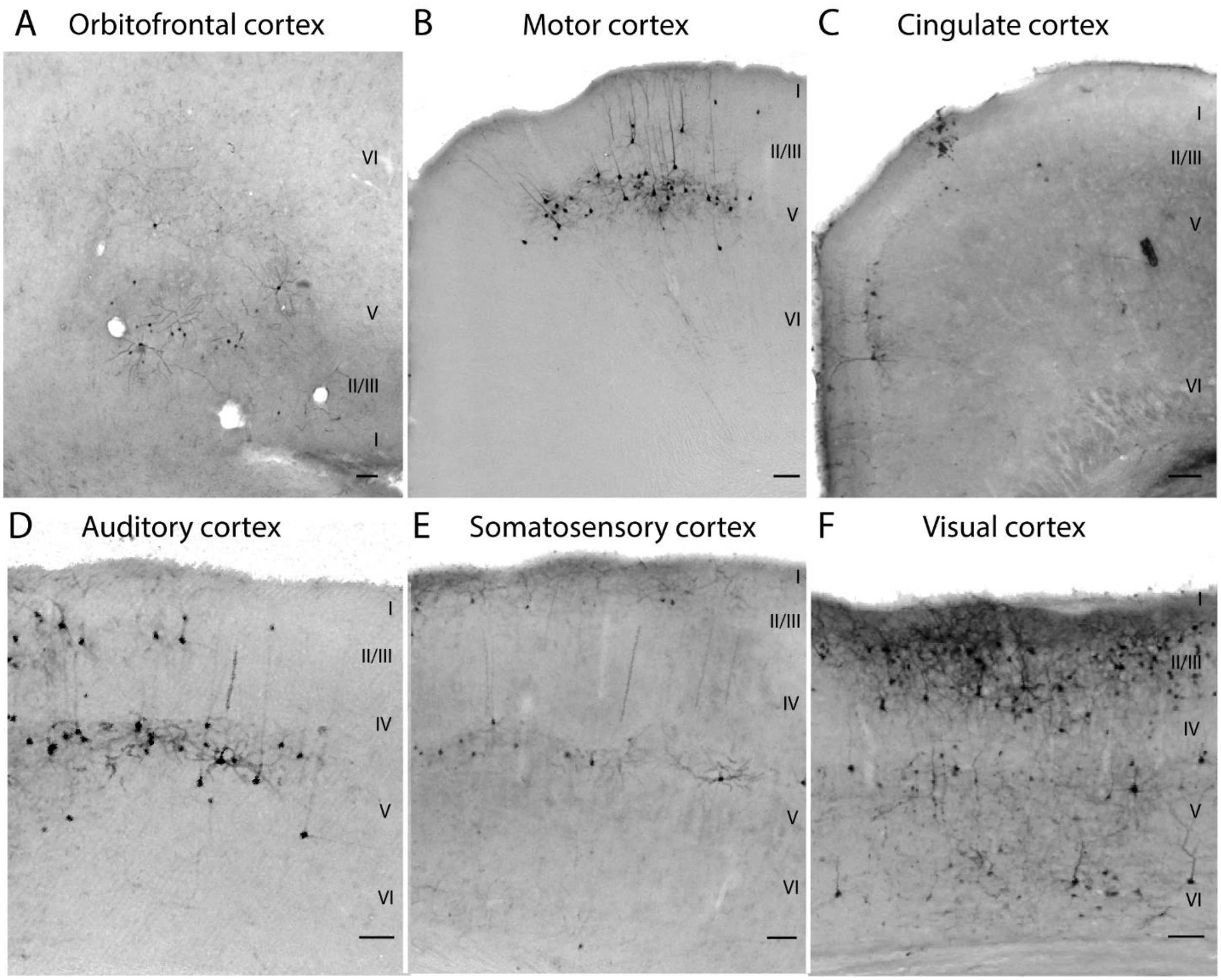


**Supplementary Figure 5 - Retrograde labeling of AM shows connectivity coming from all principle cortical areas to qualify as a PPC subarea.**

**A-F.** Example neurons projecting to AM were immunolabeled in: **A.** Orbitofrontal cortex; **B.** Secondary motor cortex; **C.** Anterior retrosplenial cortex; **D.** Auditory cortex; **E.** Somatosensory cortex; **F.** Visual cortex. Scale bar = 100µm

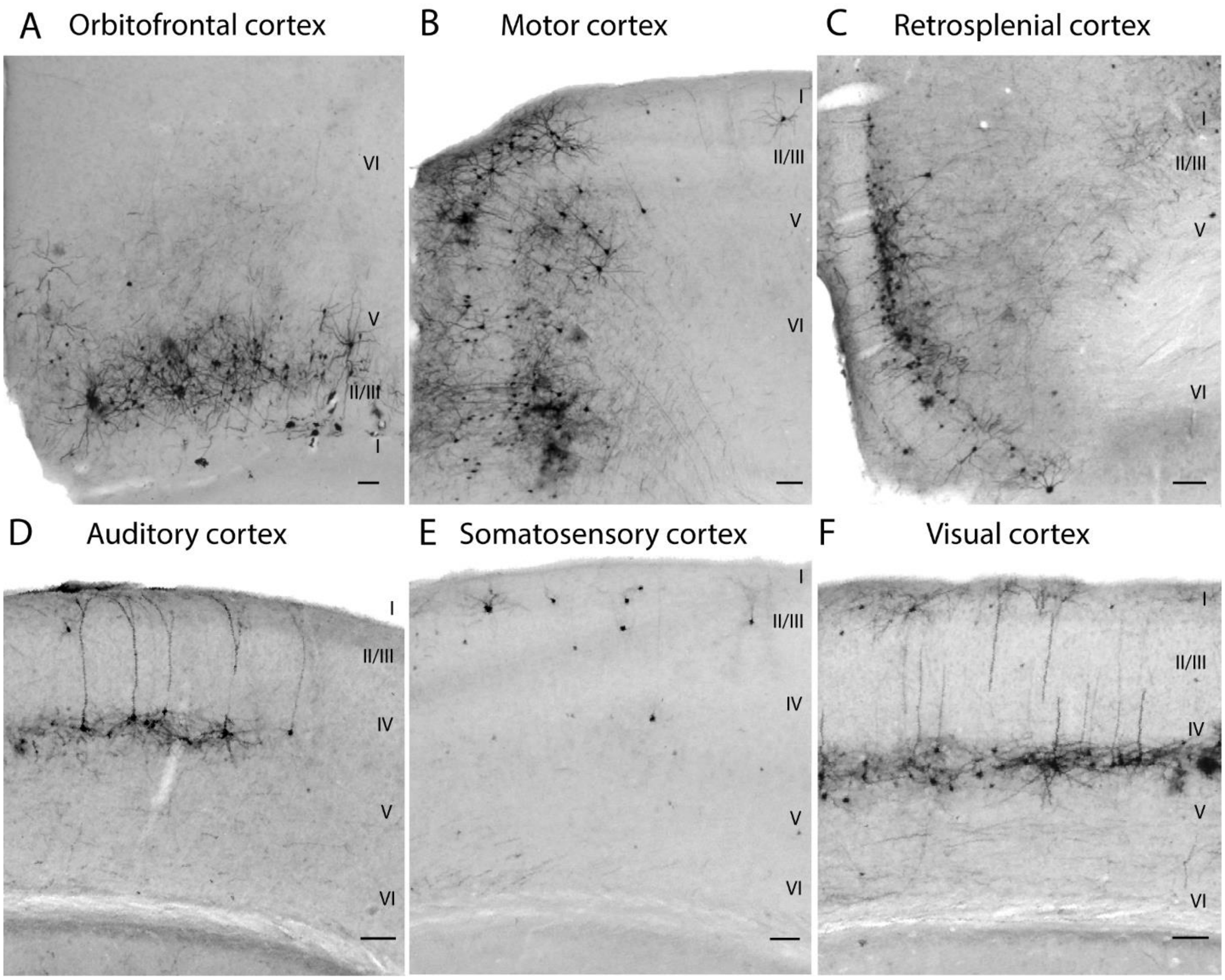


**Supplementary Figure 6 - Retrograde labeling of MMA shows connectivity coming from all principle cortical areas to qualify as a PPC subarea.**

**A-F.** Example neurons projecting to AM were immunolabeled in: **A.** Orbitofrontal cortex; **B.** Secondary motor cortex; **C.** Anterior retrosplenial cortex; **D.** Auditory cortex; **E.** Somatosensory cortex; **F.** Visual cortex. Scale bar = 100μm

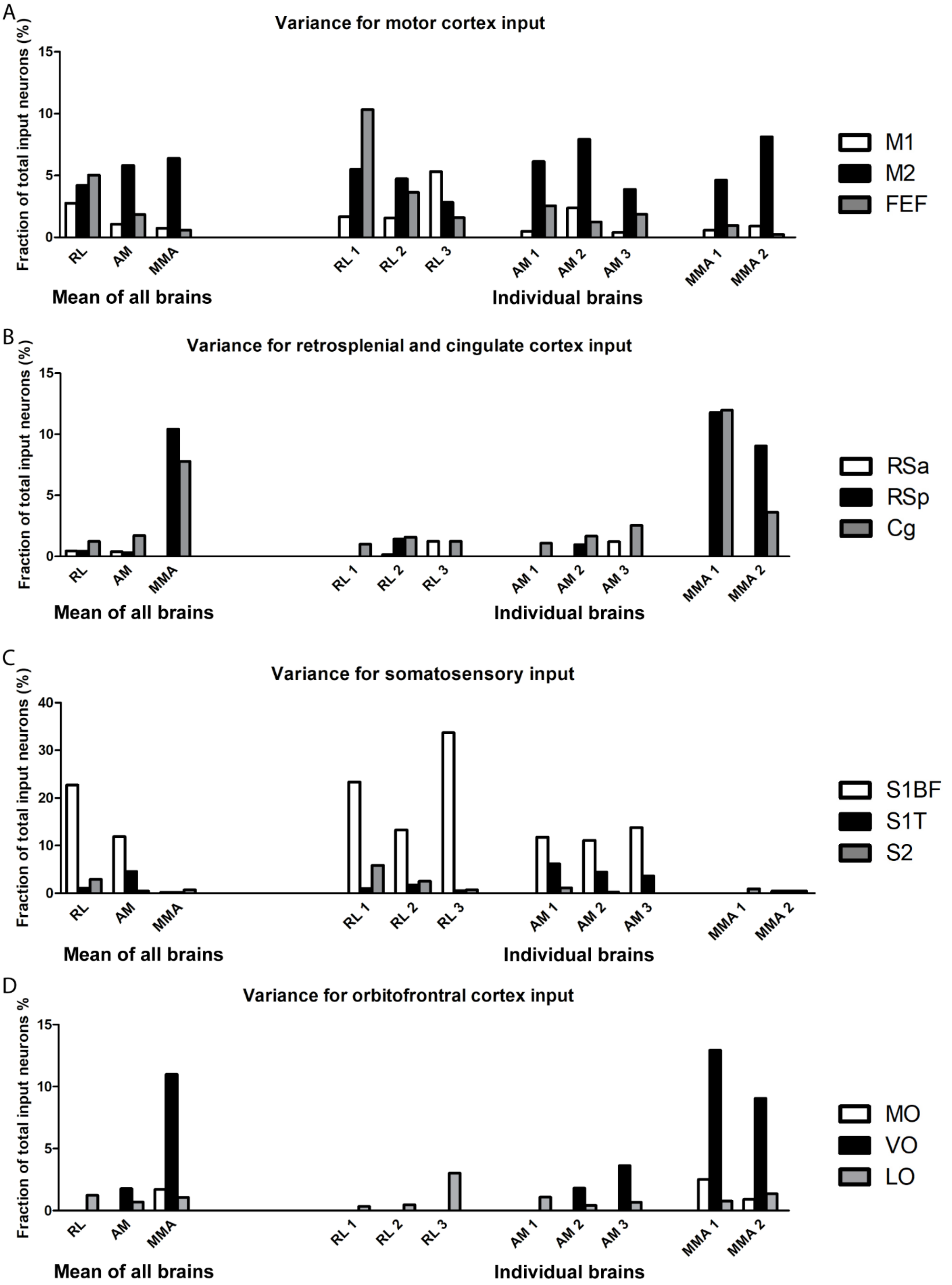
A
Variance for motor cortex input
Fraction of total input neurons (%)
0
5
10
15
RL
AM
MMA
RL 1
RL 2
RL 3
AM 1
AM 2
AM 3
MMA 1
MMA 2
Mean of all brains
Individual brains
M1
M2
FEF
B
Variance for retrosplenial and cingulate cortex input
Fraction of total input neurons (%)
RSa
RSp
Cg
C
Variance for somatosensory input
Fraction of total input neurons (%)
0
10
20
30
40
S1BF
S1T
S2
D
Variance for orbitofrontral cortex input
Fraction of total input neurons %
MO
VO
LO

**Supplementary Figure 7 – Variance in connectional weight – the fraction of labeled input neurons - between individual brains is present, yet supports the same overall conclusions.**

**A – D:** Left on the X-axis are the averaged fractions of input neurons for each area, identical to those depicted in figure 4, while the right side of the X-axis shows the fraction of input neurons for each individual brain.

**A.** Variance between individual brains for motor cortex input.

**B.** Variance between individual brains for retrosplenial and cingulate cortex input.

**C.** Variance between individual brains for somatosensory cortex input.

D. Variance between individual brains for orbitofrontal cortex input

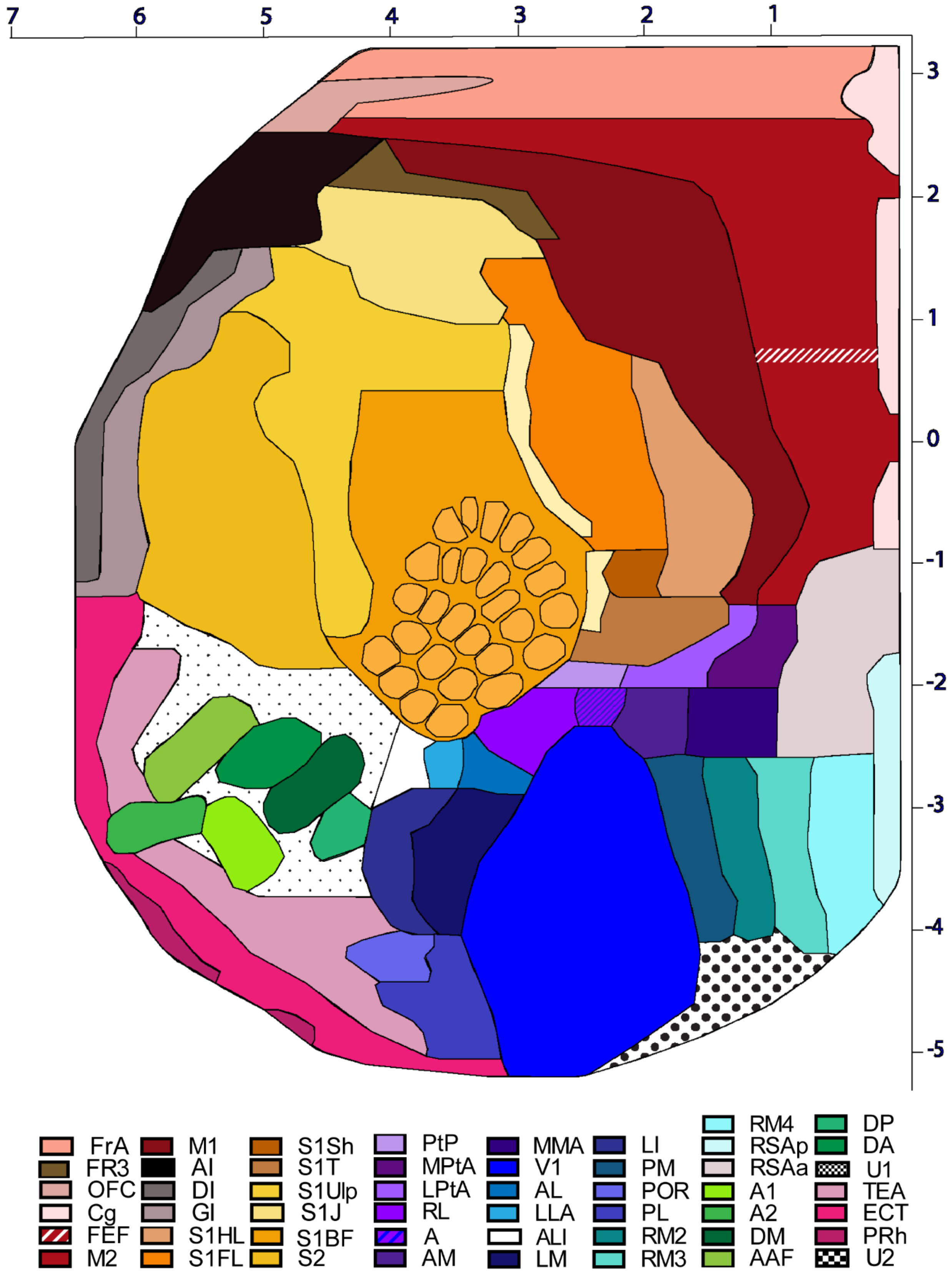
7
6
5
4
3
2
1
3
2
1
0
-1
-2
-3
-4
-5
FrA
FR3
OFC
Cg
FEF
M2
M1
AI
DI
GI
S1HL
S1FL
S1Sh
S1T
S1Ulp
S1J
S1BF
S2
PtP
MPtA
LPtA
RL
A
AM
MMA
V1
AL
LLA
ALI
LM
LI
PM
POR
PL
RM2
RM3
RM4
RSAp
RSAa
A1
A2
DM
AAF
DP
DA
U1
TEA
ECT
PRh
U2

**Supplementary Figure 8 – Large detailed flattened cortical map, representing every cortical area in a different color.**

X-axis shows the medio-lateral coordinates in mm, the Y-axis shows the antero-posterior coordinates in mm, with zero representing bregma. See list of area abbreviations in relation the area denominations.
